# Developing synergistic drug combinations to restore antibiotic sensitivity in drug-resistant *Mycobacterium tuberculosis*

**DOI:** 10.1101/860288

**Authors:** Charles Omollo, Vinayak Singh, Elizabeth Kigondu, Antonina Wasuna, Pooja Agarwal, Atica Moosa, Thomas R. Ioerger, Valerie Mizrahi, Kelly Chibale, Digby F. Warner

**Author notes:** Centre for Traditional Medicine & Drug Research, Kenya Medical Research Institute, Nairobi, Kenya (Elizabeth Kigondu).

## Abstract

Tuberculosis (TB) is a leading global cause of mortality owing to an infectious agent, accounting for almost one-third of antimicrobial resistance (AMR) deaths annually. We aimed to identify synergistic anti-TB drug combinations with the capacity to restore therapeutic efficacy against drug-resistant mutants of the causative agent, *Mycobacterium tuberculosis*. We investigated combinations containing the known translational inhibitors, spectinomycin (SPT) and fusidic acid (FA), or the phenothiazine, chlorpromazine (CPZ), which disrupts mycobacterial energy metabolism. Potentiation of whole-cell drug efficacy was observed in SPT-CPZ combinations. This effect was lost against an *M. tuberculosis* mutant lacking the major facilitator superfamily (MFS) efflux pump, Rv1258c. Notably, the SPT-CPZ combination restored SPT efficacy against an SPT-resistant mutant carrying a g1379t point mutation in *rrs*, encoding the mycobacterial 16S ribosomal RNA. Combinations of SPT with FA, which targets the mycobacterial elongation factor G, exhibited potentiating activity against wild-type *M. tuberculosis*. Moreover, this combination produced a marginal potentiating effect against both FA-monoresistant and SPT-monoresistant mutants. Finally, combining SPT with the frontline anti-TB agents, rifampicin (RIF) and isoniazid, resulted in enhanced activity *in vitro* and *ex vivo* against both drug-susceptible *M. tuberculosis* and a RIF-monoresistant *rpoB* S531L mutant.These results support the utility of novel potentiating drug combinations in restoring antibiotic susceptibility of *M. tuberculosis* strains carrying genetic resistance to any one of the partner compounds.

## INTRODUCTION

The increasing prevalence of multidrug-resistant tuberculosis (MDR-TB) – defined as resistance to the frontline anti-TB agents, isoniazid (INH) and rifampicin (RIF) – necessitates the urgent development and implementation of new anti-mycobacterial drugs and therapeutic strategies (1, 2). A number of anti-TB compounds are currently in the drug discovery pipeline, with several others in advanced preclinical development (3). However, with the exception of bedaquiline (BDQ) and delamanid, no TB-specific drugs have been introduced into clinical use within the past 40 years (4). Therefore, new options need to be explored to address the problem of drug resistance.

Novel combination regimens comprising standard anti-TB agents and repurposed drugs represent a logical approach, especially where the new partner drug has already been approved for other clinical indications (5-7). Recent advances in understanding the physicochemical properties that determine drug distributions within complex tissue and cellular (micro)environments (8, 9), together with improved methods for rapid selection of multiple potentially synergistic drug partners *in vitro* and *in vivo* (8, 10-12), suggest the potential for rational development of novel combination approaches. This is important since it might address the long-held concern that developing combinations should be avoided owing to the complexities inherent in ensuring simultaneous and sustained delivery of the optimal partner compounds to the same target site (11). The impact of pre-existing drug-resistance on the utility of new drug combinations presents an additional challenge, and is of particular concern when these combinations comprise current frontline anti-TB agents. To minimize the risks of exposing an individual to effective monotherapy, the likely pre-existence of resistance to individual drugs must be recognized, and informed combination approaches for drug therapies designed. These should incorporate multiple attributes beyond simply selecting individual molecules based on their biological activities as single agents (13).

In this study, we employed spectinomycin (SPT) as an anchor compound in combination with other experimental antibiotics and existing frontline anti-TB agents. SPT is an aminocyclitol antibiotic that inhibits protein synthesis by disrupting mRNA interactions with the 30S ribosome (14). Unlike other aminocyclitol antibiotics (including gentamycin and kanamycin), SPT is not ototoxic (15) and has been used extensively in treating *Neisseria gonorrhoeae* infections in patients who cannot tolerate first-line treatments (10). From the perspecective of new regimen design, SPT has been shown in combination screens against *M. tuberculosis* to synergize with several different classes of antimycobacterial compounds, both *in vitro* and in a macrophage model (10). Unfortunately, a key liability undermining its utility as a single agent is that SPT is subject to active efflux by *M. tuberculosis* – an observation which motivated an elegant medicinal chemistry solution in the development of the spectinamides as derivative “efflux-resistant” anti-TB antibiotics (16-18). Spectinamides are also known to synergize with a variety of antibiotic classes (11) with lead spectinamide molecules, such as 1599, shown to be active against MDR *Mtb* strains and to synergize with existing and experimental anti-TB drugs *in vivo* (19, 20). However, developing suitable spectinamide formulations for therapeutic delivery remains an obstacle to the advancement of these compounds as novel anti-TB agents (21, 22).

We investigated the utility of potentiating combinations, anchored by SPT, to circumvent drug resistance and, potentially, restore (partial) susceptibility where genetically drug-resistant mutants pre-exist for one of the partner compounds. We applied combination screens utilizing (i) chlorpromazine (CPZ), a phenothiazine whose complex and unresolved mechanism of action involving disruption of the mycobacterial electron transport chain (23) has been implicated in efflux pump inhibition (24); and (ii) fusidic acid (FA), a translational inhibitor with demonstrated (albeit moderate) activity *in vitro* (25, 26). FA was selected owing to its potential for repositioning as anti-TB agent and because it possesses a unique mechanism of action: specifically, inhibition of bacterial protein synthesis by binding to elongation factor G (EF-G) (27). The antimicrobial-potentiating effect of FA with other antibiotics, including the frontline anti-TB drug, ethambutol (EMB), as well as its lack of cross-resistance to other antimicrobial classes, provided additional motivation for our choice of FA (12, 28). By testing these combinations against both drug-susceptible *M. tuberculosis* H37Rv and selected drug-resistant mutants, we explored new potentiating combinations and demonstrated the utility of developing potent combinations against bacilli carrying pre-existing genetic resistance to either of the partner drugs. In addition, this work revealed that the inclusion of SPT as third agent with the existing first-line anti-TB drug combination of RIF and NIH restores activity *in vitro* against defined pre-MDR mutants of *M. tuberculosis*.

## MATERIALS AND METHODS

### Chemicals and reagents

All chemicals and solvents were purchased from Sigma-Aldrich. Working solutions of all antimicrobial agents were prepared in dimethyl sulfoxide (DMSO).

### Bacterial strains and culture conditions

The laboratory strain, *M. tuberculosis* H37Rv, its derivative mutants, and a reporter strain which has been used previously in high-throughput antimicrobial drug screening and constitutively expresses green fluorescent protein (GFP), H37Rv pMSP::eGFP (29), were maintained as freezer stocks. Strains were inoculated in standard Middlebrook 7H9 medium supplemented with 10% oleic acid-albumin-dextrose-catalase (OADC) (Difco) an incubated as stationary cultures at 37°C for approximately 3 days, sub-cultured, and incubated until culture density was approximatly OD_600_ ∼0.5. A second reporter mutant, *M. tuberculosis* H37Rv::(pSMYC::mCherry) (30), that constitutively expresses the mCherry fluorophore, was grown in media containing 50 mg/L hygromycin. Cell suspensions were diluted to give an expected final concentration of 10^5^ cells/ml at the time of inoculation into the microplate for the minimum inhibitory concentration (MIC) assays.

### Drug susceptibility assays

Resazurin was used to determine the susceptibility of drugs against *M. tuberculosis* strains. The MIC_90_ for all drugs was determined by broth microdilution assay in 96-well microtiter plates (31). Briefly, 2-fold serial dilutions of compounds were performed on clear, round-bottom 96-well plates using 7H9-OADC medium. *M. tuberculosis* cultures, grown to to an OD_600_ of 0.5 (∼ 10^8^ cells/ml) and diluted 1,000-fold, were added at equal volume for a total volume of 100 µl per well. The plates were sealed in zip-lock bags and incubated at 37 °C for 7 days, consistent with EUCAST guidelines (32) and published literature (33) recommending that MIC plates should be read after 7 and 14 days post inoculation. Resazurin dye was added and the plates incubated for a further 24 h. Fluorescence readings, at excitation and emission wavelengths of 540 and 590 nm respectively, were recorded using a SpectraMax i3x plate reader (Molecular Devices), and the MIC_90_, the lowest drug concentration to inhibit growth by more than 90%, was determined from the dose-response curve.

### Checkerboard assays

#### 2D Checkerboard

Standard “two-dimensional” (2D) drug-drug interactions were determined by checkerboard titration in a 96-well plate **(Fig. S1a)**. The 2D microdilution was carried out as described (34), with slight modification. Briefly, the first drug (A) to be serially diluted was dispensed (2 µl) along the x-axis (columns 3 to 11, row B) at a starting concentration 100 times higher than the final concentration in the well, and (2 µl) per well of the second drug (B) was serially dispensed along the y-axis (from row B to H) at a starting concentration 100 times higher than the final concentration in the 96-well microtitre plate. The first column (column 1) and last column (column 12) contained drug-free controls (with 1% DMSO as a diluent) and a control drug concentration giving maximum inhibition, respectively. The second column from B2-H2 and first row from A3-A11 contained individual drugs, thus providing the MIC for each drug alone in each assay (each plate). The plates were placed in zip-lock bags and incubated at 37 °C for 7 days. Resazurin dye was then added and the plates incubated for a further 24 h. A visible colour change from blue to pink indicated growth of bacteria, and the visualized MIC was defined as the lowest concentration of drug that prevented growth (at which the colour change did not occur) (31). Fluorescence readings (excitation: 544 nm, emission: 590 nm) were obtained using BMG Labtech POLARstar Omega plate reader (BMG Labtech, Offenburg, Germany). The mean fluorescence value for the “maximum inhibition” wells (column 12) was subtracted from all other wells to control for background fluorescence. Percent inhibition was defined as 1-(test well fluorescence units/mean fluorescence units of maximum inhibition wells) × 100 on day 8 after incubation. The lowest drug concentration effecting inhibition of 90% was considered the MIC_90_ In addidition, synergy was interpreted according to the sum of fractional inhibitory concentration (ΣFIC). The fractional inhibitory concentration for each compound was calculated as follows (35): FIC_A_ _(MIC of compound A in the presence of compound B)/(MIC of compound A alone), where FIC_A_ is the fractional inhibitory concentration of compound A. Similarly, the FIC for compound B was calculated. The ΣFIC was calculated as FIC_A_ + FIC_B_. Synergy was defined by values of ΣFIC ≤ 0.5, antagonism by ΣFIC > 4.0, and no interaction by ΣFIC values from 0.5 to 4.0 (36).

#### 3D Checkerboard

In the three-drug (“three-dimensional”, 3D) combinations **(Fig. S1b)**, microdilutions for the first two drugs were initially set up principally following the standard 2D checkerboard assay protocol described above. The third drug (2 µl) was then added at a starting concentration 100 times higher than the final concentration in the well as an overlay at five sub-inhibitory concentrations ranging from 1/32 to 1/2 of the single-drug MIC. Well A2 on all plates contained the third drug only, providing the single-drug MIC for the third drug in each assay (set of 5 plates). After inoculation with a log-phase culture, OD_600_ of 0.5 (∼ 10^8^ cells/ml) of *M. tuberculosis*, 50 µl to each well, the plates were placed in zip-lock bags and incubated for 7 days at 37°C before addition of resazurin. The plates were further incubated for 24 h and the results were read in the BMG Labtech plate reader (excitation 544 nm; emission 590 nm) (33). The % inhibition was calculated as described earlier.

### Macrophage assays

#### Cell culture and maintenance

Human promonocytic THP-1 cells were maintained in RPMI-1640 medium (Sigma) supplemented with 10% fetal bovine serum (FBS, Invitrogen) at an initial density of 8×10^5^ cells/ml at 37°C in an humidified, 5% CO_2_ atmosphere. Prior to plating of the cells, viability was assessed by trypan blue exclusion method (37). Maturation of THP-1 cells into macrophages was induced by adding 200 nM PMA (phorbol 12-myristate 13-acetate; Sigma) in cell culture medium for 24 h. Differentiated macrophages were then washed three times with pre-warmed phosphate buffer saline (PBS) to remove the PMA and replenished with cell culture medium.

#### Infection of macrophages and drug treatment

To check the efficacy of drugs in macrophages, 5×10^4^ THP-1 cells/well (100 µl final volume) in 96-well flat-bottomed tissue culture plates were differentiated into macrophages. To infect macrophages, an exponentially growing *M. tuberculosis* H37Rv::(pSMYC::mCherry) culture was harvested by centrifugation and washed twice with PBS. The pellet was resuspended in PBS and passed through a 5 µm filter to generate a suspension of single-cell bacilli. The bacterial suspension density was estimated by measuring OD at 600 nm, corresponding an OD_600_∼0.5 to 1×10^8^ CFU/ml. Infection medium comprised cell culture medium containing a number of bacteria required to achieve a multiplicity of infection (MOI) of 5:1 (5 bacilli for every THP-1 cell) (38). The macrophage cells were overlaid with infection medium and incubated at 37°C in 5% CO_2_ for the phagocytic period of 3 hours. Cells were then washed gently and thoroughly with prewarmed PBS to remove extracellular bacteria. The cells from three wells were lysed by adding triton X-100 (0.05% in PBS) and the lysate was plated onto 7H10 to score colony forming units (CFUs) for untreated day zero. The used as untreated control were refreshed with cell culture medium. The cells in remaining wells were treated with the indicated antibiotic either alone or in combination at 1× MIC_90_ or 5× MIC_90_, as determined in liquid culture. Hygromycin at 50 mg/L was added into the cell culture medium for all wells to maintain the plasmid expressing mCherry.

#### Fluorescence measurement

The fluorescence of the mCherry reporter (excitation: 590 nm, emission: 610 nm) was measured at different time points on a BMG Labtech plate reader.

#### CFU enumeration

To estimate the numbers of live bacilli after drug treatment, untreated and drug-treated *M. tuberculosis*-infected cells were lysed in triton X-100 (0.05% in PBS) on days 2, 4 and 6, and serial dilutions of the cell lysate were plated onto 7H10 agar. Colonies were counted after 3-4 weeks of incubation at 37 °C.

### Statistical Analyses

Statistical analyses were performed using Prism 9.0.0.121 (GraphPad). Means were compared via ANOVA, with post-test evaluation using Dunnett’s or Bonferroni’s test. *P* values are abbreviated as follows: *, *P*<0.05; **, *P*<0.01; \*\*\**P*<0.001

## RESULTS

### CPZ potentiates SPT activity by inhibiting Rv1258c-mediated efflux

The combination of SPT and CPZ was previously reported as synergistic against *M. smegmatis* (39). When tested against wild-type *M. tuberculosis* H37Rv **(Fig. 1a)**, the same combination yielded a ΣFIC value of 0.09 **(Table 1)**, confirming strong synergy (36). We investigated whether this effect resulted from CPZ-mediated disruption of the activity of the Major Facilitator Superfamily (MFS) efflux pump, Rv1258c, which has been implicated in innate resistance to SPT (17). We performed checkerboard assays using the efflux-defective Δ*Rv1258c* (“tap”) knockout mutant, which had been used in the development of the spectinamides (SPD) (17), and its complemented derivative, Δ*Rv1258c* pCRS4. Both mutants exhibited the same MIC_90_ of 22 mg/L for CPZ **(Fig. 1** & **Table 1)**, whereas the Δ*Rv1258c* knockout mutant was hypersusceptible to SPT, returning an approximately 6-fold lower MIC_90_ of 3.9 mg/L **(Fig. 1b, Table 1 and Table S1)**, as observed previously (40). Notably, the synergy detected on exposing wild-type *M. tuberculosis* to a combination of CPZ and SPT (**Fig. 1a**) was eliminated in the Δ*Rv1258c* mutant **(Fig. 1b)** – which yielded a ΣFIC value of 0.75 **(Table 1)** – but was restored in the complemented Δ*Rv1258c* pCRS4 strain **(Fig. 1c)**, with a ΣFIC of 0.12 **(Table 1)**. Previous studies have reported no significant alteration in *Rv1258c* transcription in response to CPZ treatment (41). Therefore, our observations suggested that CPZ treatment abrogated efflux-mediated intrinsic resistance to SPT in wild-type *M. tuberculosis* in a manner dependent on Rv1258c, perhaps as a result of CPZ-mediated inhibition of energy metabolism (23).

**Table 1.**
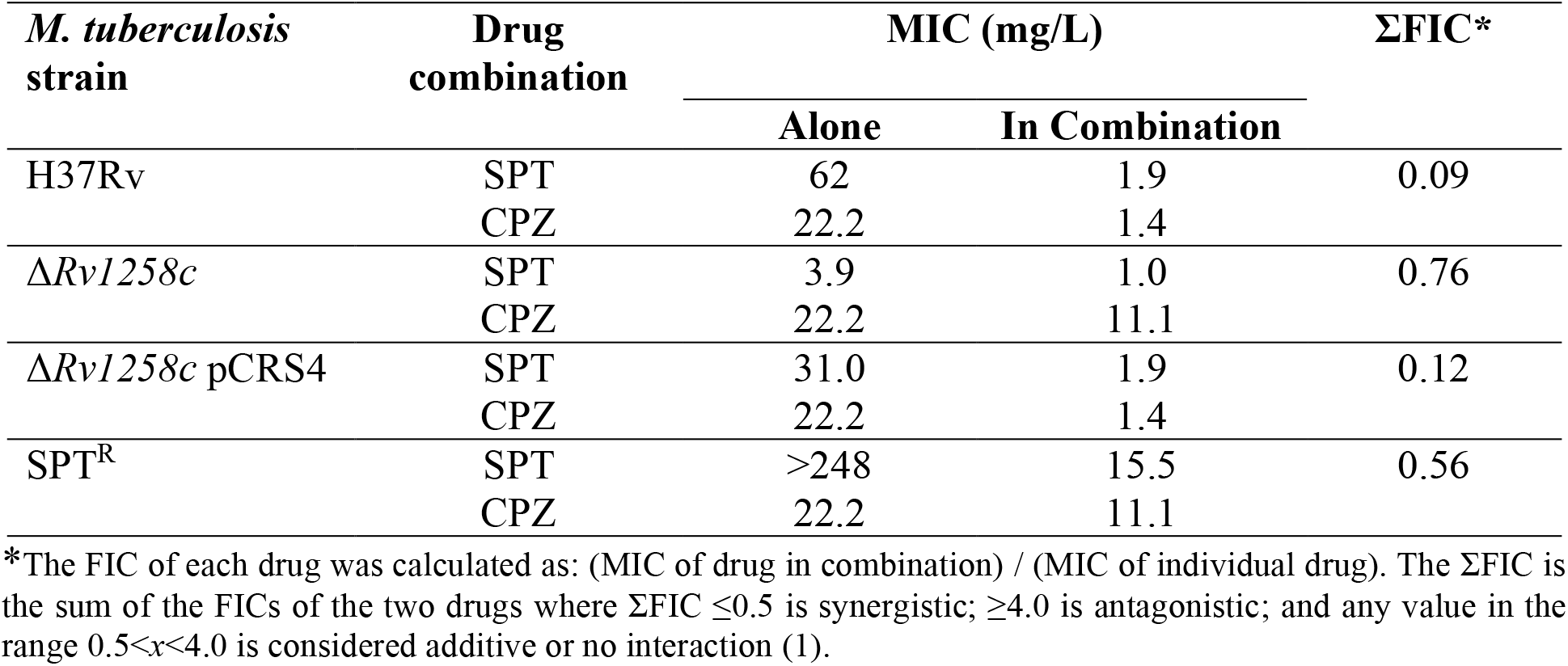
Investigation of potential synergies between SPT and CPZ against different *M. tuberculosis* strains through the calculation of the FIC and sum of the FIC.

**Figure 1.**
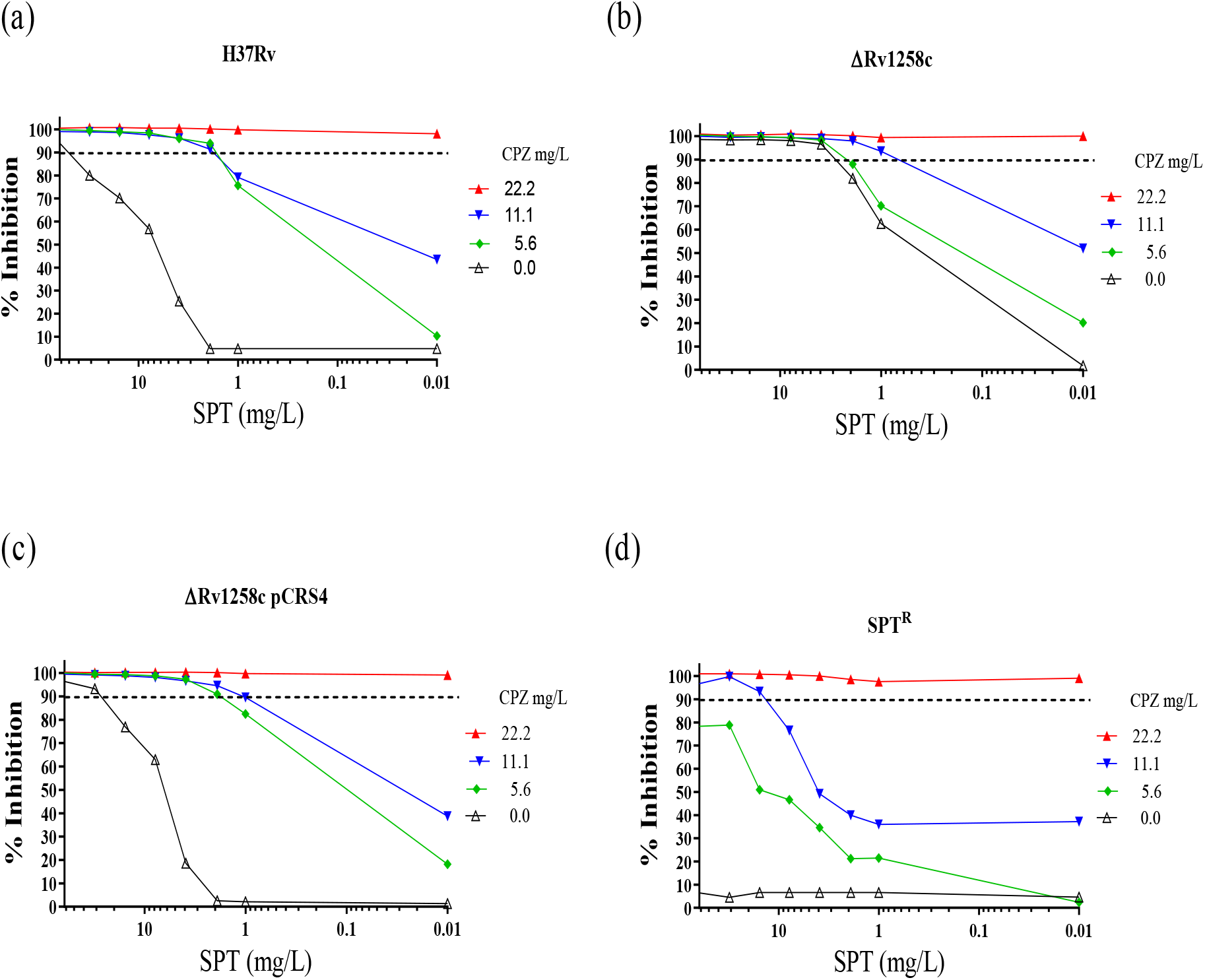
Inhibition of Rv1258c-mediated efflux of SPT by addition of CPZ. Combinations of CPZ and SPT were applied in checkerboard assays against (*a*) wild-type *M. tuberculosis* H37Rv, (*b*) the Δ*Rv1258c* deletion mutant, (*c*) the Δ*Rv1258c* pCRS4 complemented mutant, and (*d*) the SPT^R^ strain. Bacterial growth inhibition was assessed in two independent experiments by fluorescence-based resazurin assay. The dashed horizontal line indicates 90% inhibition.

### CPZ synergizes with compounds other than SPT, but SPT potentiation arises solely from CPZ-mediated inhibition of Rv1258c

To determine whether the synergy observed was specific for SPT or might apply to other antimycobacterial agents, CPZ was applied as the anchor compound in combination assays with a panel of anti-TB antibiotics of different classes and mechanisms of action (**Table S2**). Of the eight compounds tested with CPZ, four exhibited clear synergy (ΣFIC ≤ 0.5): the frontline agents, RIF and INH, which returned ΣFICs of 0.37 and 0.5, respectively, and bedaquiline (BDQ) and nalidixic acid, both of which gave ΣFICs ≤ 0.25. In contrast, no potentiation was observed with KAN or the fluoroquinolones, ciprofloxacin (CIP) and levofloxacin (LEV), all of which yielded ΣFICs > 0.5.

In a complementary approach, we also investigated if the potentiating effect observed with the SPT-CPZ combination was unique to CPZ. To this end, we assayed SPT in combination with an expanded panel of antimycobacterial agents (**Fig. S2** and **Table S3**). SPT was found to synergize with only two of the eleven compounds tested: erythromycin (ERY), a macrolide targeting the ribosome, and verapamil (VER), a cationic amphiphile which was originally considered an *M. tuberculosis* efflux pump inhibitor but has been shown recently to disrupt membrane function (42). RIF, the mycobacterial RNA polymerase inhibitor, was just beyond the threshold determining synergistic activity.

To ascertain if inhibition of Rv1258c-mediated efflux resulted in the observed compound synergies, the Δ*Rv1258c* mutant was tested for hypersensitivity to a corresponding panel of anti-TB agents **(Fig. S3** and **Table S1)**. The spectinamide 1599 had the same MIC_90_ for both wild-type *M. tuberculosis* H37Rv and the Rv1258c mutant, reflecting its successful modification to avoid Rv1258c-mediated efflux (43). Of the other 11 compounds tested, only SPT was associated with hypersensitivity in the Rv1258c knock-out mutant, returning an MIC_90_ value of 0.39 mg/L compared to the MIC_90_ value of 62-125 mg/L against wild-type strain. In combination, these results strongly support the inference that the synergy detected with the CPZ-SPT combination arises from CPZ-mediated inhibition of Rv1258c.

### The CPZ-SPT combination partially restores SPT sensitivity in an SPT-resistant mutant

A spontaneous SPT-resistant mutant (SPT^R^) carrying a g1379t point mutation in the mycobacterial 16S ribosomal RNA, *rrs*, was associated with >64-fold increase in the SPT MIC_90_ **(Table S1)**. In contrast, the activity of CPZ remained at the wild-type concentration for this strain, consistent with a mechanism of action of CPZ that was independent of *rrs* inhibition (44). While the SPT^R^ mutant was resistant to SPT at concentrations >248 mg/L in the absence of CPZ, combinations utilizing CPZ at sub-MIC concentrations ([CPZ] ≤ 11.1 mg/L) restored SPT sensitivity, at least partially **(Fig. 1d)**. These results suggested the capacity for synergistic combinations to restore drug activity against mutant strains genetically resistant to either of the partner compounds.

### Assessing synergy with SPT and fusidic acid, two antibiotics acting on the mycobacterial translational machinery

The systematic application of drug combinations can reveal synergistic interactions. One form of synergy occurs when drug(s) which perturb normal cell physiology trigger (compensatory) cellular responses that can, in turn, affect (potentiate) the activities of other drugs (45). Nichols *et al*. demonstrated that the synergistic interaction between sulfamethoxazole and trimethoprim was a result of the two drugs targeting tetrahydrofolate biosynthesis (46). With this in mind, a combination comprising SPT and FA – translation inhibitors which act at discrete steps of the elongation process – was tested against wild-type *M. tuberculosis* and two resistant strains, FA^R^ and SPT^R^. Isolation of spontaneous FA^R^ mutants *in vitro* yielded a single strain on 25× MIC FA at a frequency ∼10^−8^. Whole-genome sequencing identified a c1384t (H462Y) substitution in *fusA1* (*Rv0684*), encoding the essential mycobacterial elongation factor G (EF-G) (48). The histidine residue is highly conserved across multiple bacterial species; therefore, using the *T. thermophilus* structure as template (49), it can be inferred that *M. tuberculosis* H462 corresponds to *T. thermophilus* H458 (50), mutations of which are likely to alter the FA-binding pocket (51). In MIC assays, the H462Y mutant consistently yielded an MIC_90_ ≥ 25 mg/L, confirming heritable FA^R^ **(Table 2)**.

**Table 2.** Investigation of potential synergies between SPT and FA against different *M. tuberculosis* strains through the calculation of the FIC and sum of the FIC

| *M. tuberculosis*<br>strain | Drug<br>combination | MIC (mg/L) | | ΣFIC |
| --- | --- | --- | --- | --- |
|  |  | Alone | In Combination |  |
| H37Rv | SPT | 62 | 16.3 | 0.5 |
|  | FA | 0.63 | 0.16 |  |
| SPT<sup>R</sup> | SPT | ≥62 | ≥62 | 1.5 |
|  | FA | 0.63 | 0.32 |  |
| FA<sup>R</sup> | SPT | 62 | 31.5 | 1.5 |
|  | FA | ≥25 | ≥25 |  |

\*The FIC of each drug was calculated as: (MIC of drug in combination) / (MIC of individual drug). The ΣFIC is the sum of the FICs of the two drugs where ΣFIC ≤0.5 is synergistic; ≥4.0 is antagonistic; and any value in the range 0.5< $\Sigma$ <4.0 is considered additive or no interaction (36).

With these strains in hand, we evaluated the interaction between SPT and FA and, furthermore, assessed whether this combination – which is synergistic against the parental, drug-susceptible *M. tuberculosis* H37Rv – might counter pre-existing genentic resistance to either compound. The combination of SPT and FA returned a ΣFIC value of 0.50 **(Table 2)** against wild-type H37Rv; upon addition of FA at sub-MIC concentration ([FA] = 0.16 mg/L), the MIC_90_ of SPT exhibited a ∼4-fold decrease from 62 mg/L to 16.3 mg/L **(Fig. 2a)**, p<0.001. FA at sub-inhibitory concentration ([FA] = 0.32 and [FA] = 0.16 mg/L) enhanced the activity of SPT ([SPT] = 62 mg/L) against a SPT^R^ mutant. The inhibitory effect was significantly different from the results observed with similar concentrations of SPT in the absence of FA ([FA] = 0 mg/L), p<0.001 **(Fig. 2b)**. Although the calculated sum FIC did not satisfy the criterion for “synergistic” (ΣFIC≤0.5) **(Table 2)**, the effect was marked and reproducible in two independent biological replicates **(Fig. 2b)**. Notably, the same combination did not return enhanced activity against the FA^R^ mutant **(Fig 2c)**, strongly suggesting that FA was the major contributor to the SPT-FA combination. A summary of the inhibitory effects of the CPZ-SPT and FA-SPT combinations is presented in **Table 3**.

**Table 3.**
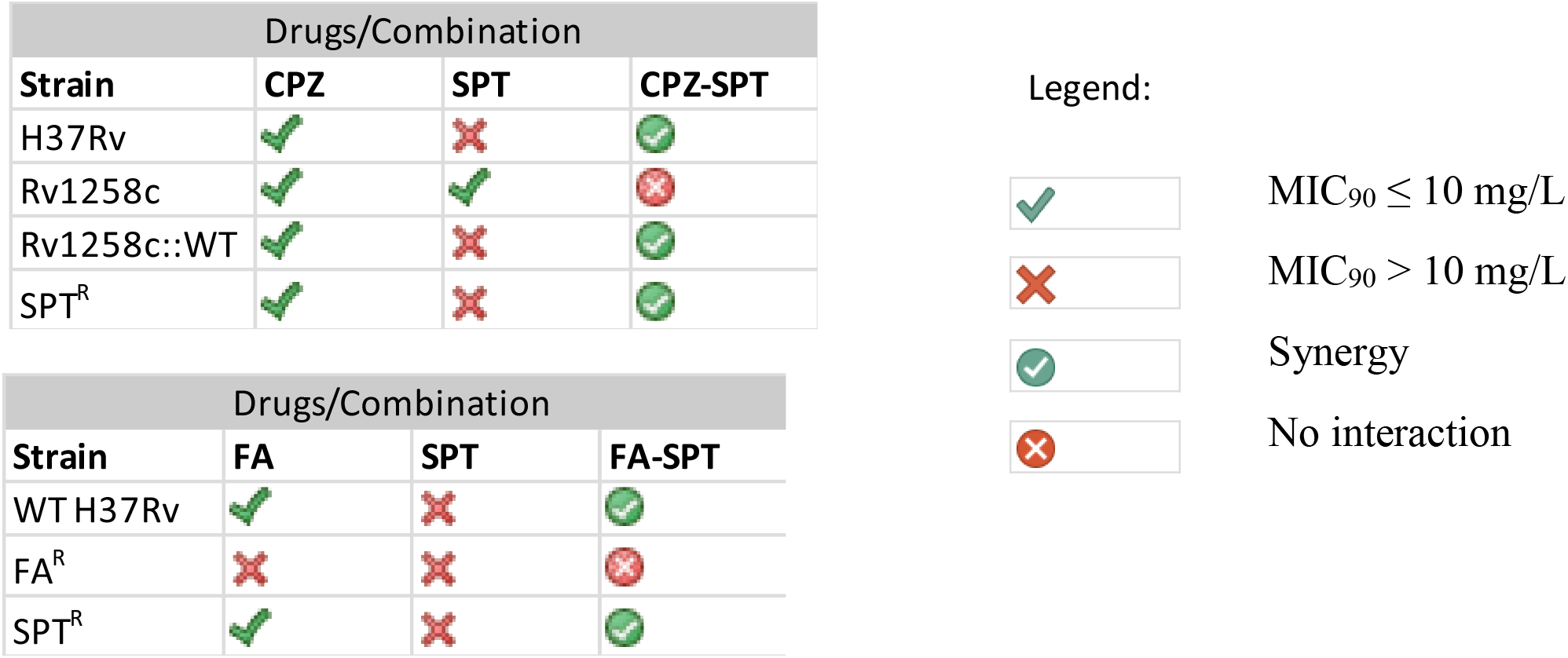
Summary of *in vitro* drug activities against *M. tuberculosis* strains

**Figure 2.**
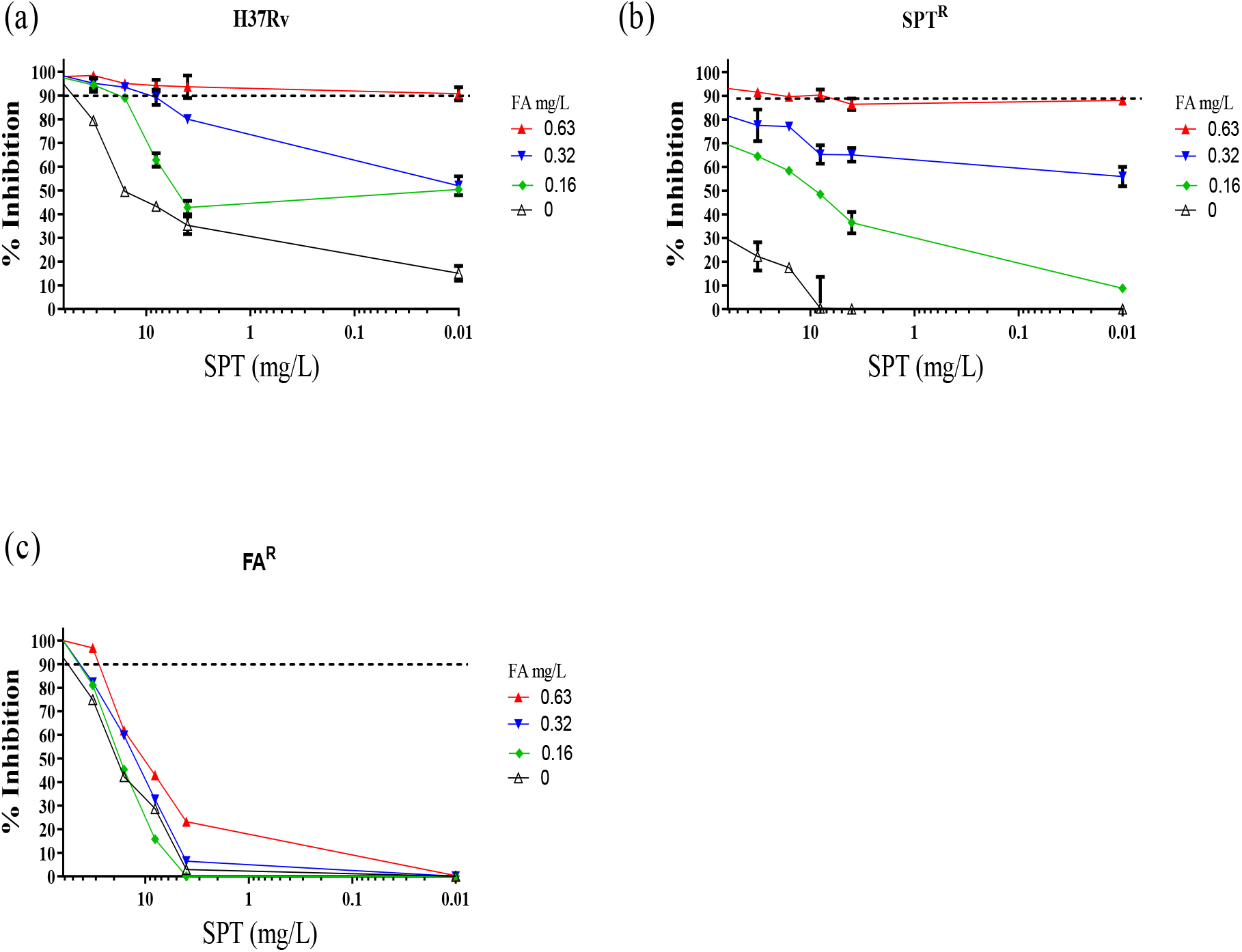
*In vitro* interaction between SPT and FA. Combinations of SPT and FA were applied in checkerboard assays against (*a*) wild-type *M. tuberculosis* H37Rv, (*b*) the SPT^R^ mutant and (*c*) the FA^R^ mutant. Bacterial viability was assessed by fluorescence-based resazurin assay. Dashed horizontal lines indicate 90% inhibition and data are the means and standard deviations of two independent biological replicates.

### Confirmation that fluorescence intensities correlate with cell density readings

The centrality of the resazurin microtiter assay (REMA) in determining the synergies reported in this study demanded orthogonal evidence supporting the claimed results. To this end, a 96-well based antimycobacterial assay was performed with a selected panel of drugs having different mechanisms of action **(Fig. S4)**. The experiments were conducted using the H37Rv::GFP reporter mutant in which expression of the fluorophore is constitutive (29). After 8 days of incubation in the presence of 2-fold dilutions of the antimicrobial agents, GFP and resazurin fluorescence intensities were determined, and the corresponding optical density read-outs recorded in parallel. Strong agreement was discerned when comparing GFP and OD_600nm_ methods with the standard resazurin read-out **(Fig. S4a & b)**, supporting the use of the resazurin assay for both MIC and FIC determinations.

### A three-drug combination comprising SPT, RIF and INH enhances *in vitro* activity against *M. tuberculosis*

The premise that synergistic combinations might be usefully applied to overcome existing drug resistance was further explored using SPT in combination with the frontline anti-TB agents, RIF and INH. In a 2D pairwise screening assay, we evaluated two-drug permutations of RIF, INH and SPT. The RIF-INH combination showed no interaction **(Fig. S5a)** while sub-MIC concentrations of INH (0.125 mg/L) and RIF (0.003 mg/L) halved the effective SPT concentration from 125 mg/L to 62 mg/L **(Fig S5b & S5c)**. To leverage the potential effect of SPT, a 3D combination assay was performed in which RIF and INH were titrated against decreasing sub-MIC_90_ concentrations of SPT (1/2×, 1/4×, 1/8×, 1/16× and 1/32× MIC_90_), using the format illustrated in **Fig. S1b**. When tested against drug-susceptible *M. tuberculosis* H37Rv, RIF at sub-MIC ([RIF] = 0.003 mg/L) plus SPT at both 1/4× and 1/2× MIC resulted in an 8-fold decrease in the effective concentration of INH from 0.25 mg/L to 0.03 mg/L **(Fig. S6)**. A kill kinetics assay showed that the addition of SPT to the RIF-INH combination elicited ∼1 log_10_ unit reduction (p < 0.001) in the viable bacillary population following 8-day exposure to the three-drug combination **(Fig. S7a)**. In contrast, when the Δ*Rv1258c* “tap” knock-out mutant was tested, RIF at sub-MIC ([RIF] = 0.003 mg/L) plus 1/2× MIC SPT resulted in only a ∼3-fold decrease in the effective concentration of INH – from 0.25 mg/L to 0.09 mg/L **(Fig. S7b)**, again implicating Rv1258c in intrinsic antibiotic resistance in *M. tuberculosis*.

### The RIF-INH-SPT combination is active in *M. tuberculosis*-infected macrophages and against monoresistant pre-MDR strains

Since *M. tuberculosis* survives and replicates in macrophages (52), the synergy of RIF-INH plus SPT was evaluated against intracellular bacilli in *M. tuberculosis*-infected THP-1 cells. This three-drug combination showed inhibitory activity at 1× MIC_90_ of the combined drugs **(Fig. S8)**. In contrast, the inhibitory effect was reduced when similar concentrations of each drug were applied individually, or when the standard RIF-INH combination was used without SPT. The intracellular activity of this triple combination was further confirmed by CFU enumeration **(Fig. S9)**, which revealed a 2-log_10_ decrease in CFU/ml when 1× MIC_90_ RIF-INH-SPT was applied compared to the untreated control.

To evaluate the efficacy of RIF-INH-SPT against known drug-resistant strains, the combination was tested against two pre-MDR *M. tuberculosis* mutants: a RIF-monoresistant mutant carrying the common *rpoB* S531L allele, and an INH-monoresistant strain harbouring a - c15t mutation in the *inhA* promoter region that confers low-level INH resistance **(Fig. 3)**. Duplicate checkerboard experiments showed that, for the *rpoB*^S531L^ mutant, addition of 1/2× MIC SPT to the RIF-INH plate resulted in a decrease in the effective concentrations of both RIF (20 to 10 mg/L) and INH (0.25 to 0.03 mg/L) **(Fig. 3a** and **3b)**, indicating partial restoration of drug susceptibility in the presence of SPT. Enhanced susceptibility was also observed for the INH^R^ mutant, albeit to a lesser extent: a sub-MIC RIF concentration of 0.01 mg/L and INH at 2.5 mg/L achieved ∼80 % bacterial inhibition **(Fig. 3c and 3d)**.

**Figure 3.**
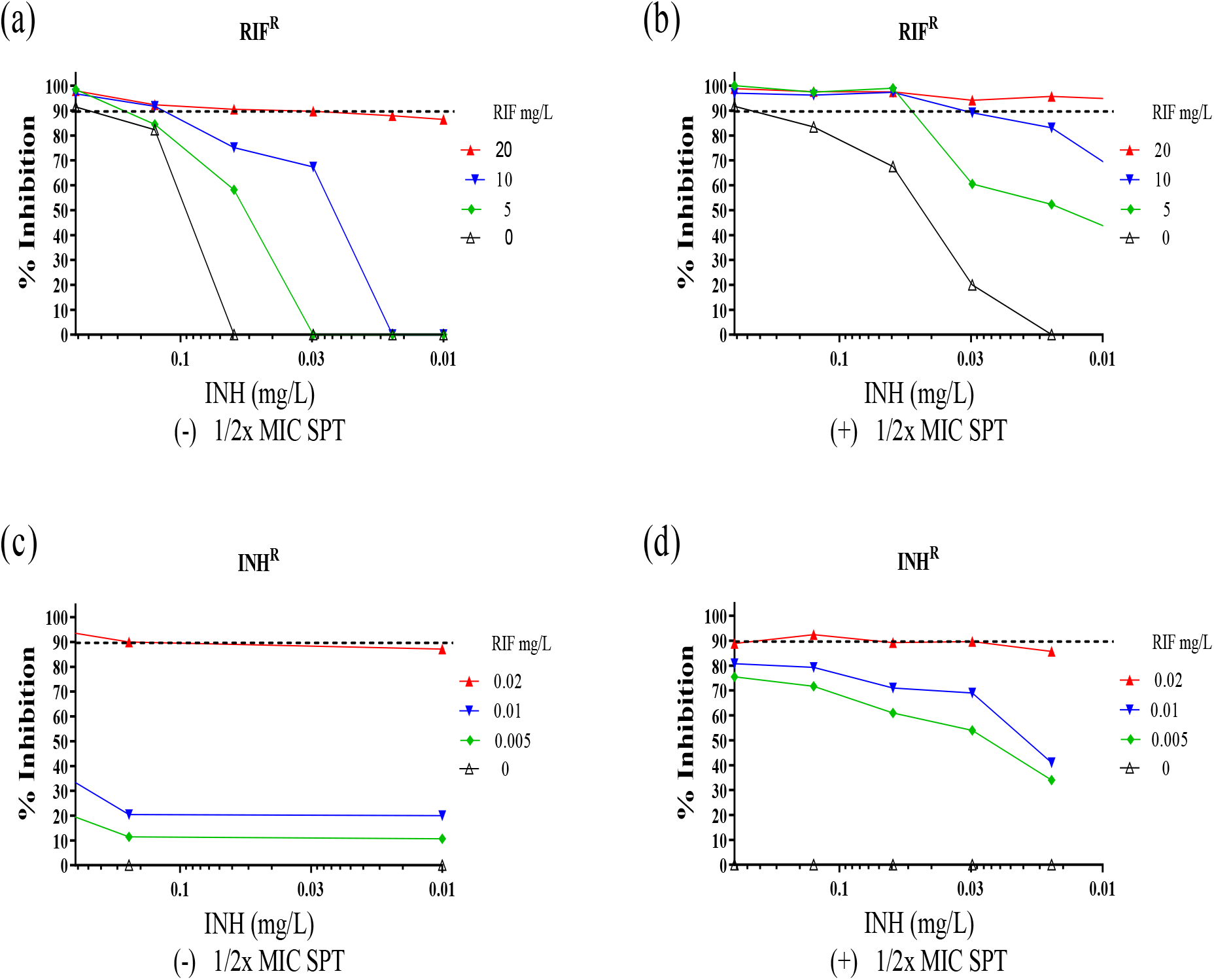
Activity against pre-MDR *M. tuberculosis* strains. *In vitro* activity of RIF-INH against RIF^R^ *M. tuberculosis rpoB*^S531L^ mutant in the (*a*) absence or (*b*) presence of 1/2× MIC SPT, and against the INH^R^ *M. tuberculosis inhA* mutant in the (*c*) absence or (*d*) presence of 1/2× MIC SPT. Bacterial viability was assessed in two independent experiments by fluorescence-based resazurin assay. The dashed horizontal line indicates 90% inhibition.

## DISCUSSION

Notwithstanding recent promising claims (53), the bacterial capacity for acquisition of resistance by multiple mechanisms means that it is difficult, perhaps even conceivably impossible, to overcome antibiotic resistance sustainably (54). Different approaches can be used to circumvent resistance transiently, ensuring antibiotic efficacy despite the pre-existence of resistant organisms in an infecting population. Combination therapy represents one such approach.

In a previous study, Chen *et al*. demonstrated a synergistic interaction between SQ109 and RIF when tested against RIF^R^ isolates: at 0.5× MIC, SQ109 was able to increase RIF’s activity against *de facto* resistant organisms in a dose-dependent manner (34). Recently, Yang *et al* reported the enhanced efficacy of the imipenem-colistin combination against multiple drug-resistant *Enterobacter cloacae in vitro* and in an infection model (55). Relativly few studies have been undertaken to illustrate the association between potentiating drug interactions and the ability of the particular drug combination to overcome pre-existing genetic resistance. In a clinical study, Ankomah *et al*. suggested that drugs acting synergistically can prevent treatment failure even when bacteria resistant to one of these drugs are present at the beginning of therapy (56). Our interaction studies between SPT and CPZ reaffirmed the susceptibility of SPT to Rv1258c-mediated efflux, an observation which suggests that efforts to modify SPT – including through novel chemical modifications that engineer resistance to efflux (17) – should be pursued.

We tested the susceptibility of the Δ*Rv1258c* mutant to a small panel of anti-mycobacterial compounds with different mechanisms of action and observed that SPT alone was associated with hypersusceptibility. A similar hypersusceptibility phenotype was achieved against wild-type *M. tuberculosis* H37Rv via chemical potentiation of SPT using CPZ as the combination agent. However, the potentiating effect of CPZ was not specific to SPT and was instead observed for a handful of other agents. This suggests the likelihood that CPZ might disable more than one intrinsic resistance mechanism in *M. tuberculosis*. Further work is required to ascertain the precise mechanism for each compound, with evidence to date implicating multiple potential efflux systems in intrinsic resistance to INH, RIF, and the fluoroquinolones (57). For BDQ, the multi-substrate RND-family transporter, MmpS5-MmpL5, appears a strong candidate based on previous reports (58).

Our results revealed synergy between FA and SPT against drug-susceptible bacteria via a mechanism independent of the efflux inhibition seen with SPT-CPZ. Notably, the same FA-SPT combination exerted an enhanced inhibitory effect against a genotypically confirmed SPT^R^ mutant compared to a FA^R^ mutant. Although there is no definitive explanation for this finding, we postulate that the relative potency of FA (∼0.63 mg/L) against the drug-susceptible H37Rv parent compared to that of SPT (∼50 mg/L) could impact the FA-SPT combination against the SPT^R^ mutant, restoring susceptibility. A similar effect was not evident in a SPT-FA combination against FA^R^ mutant owing to the diminished activity of FA and high MIC_90_ of SPT. Of interest is the impact of individual active drugs in driving synergy.

Other explanations for potentiation include the sustained drug pressure emanating from the drug interactions which leads to an increased effective dose of the drug combination. Moreover, some studies have demonstrated that the drug susceptibility of pathogens can be significantly enhanced as a result of a reduced efflux pump efficiency either by genetic manipulation (59), or addition of efflux pump inhibitors (60, 61). The clinical relevance of this finding is that, despite the existence of bacterial resistance against a combination partner, it would still be possible to achieve optimal therapeutic outcomes via the use of appropriate potent drug combinations.

Previous work has demonstrated the potential of having three-drug combinations when compared to individual or two-drug regimens (62, 63). Recently, Tekin *et al*. reported that combinations of three different antibiotics can often overcome antimicrobial resistance to antibiotics, even when none of the three antibiotics on their own — or even two of the three together — is effective (64). In addition, based on drug interaction studies, Ramon-Garcia *et al*. hypothesized that the synergistic activity of the triplet combination might have multiplicative effects (10). Here, SPT was deployed as part of a three-drug regimen which also included RIF and INH, the two drugs that form the cornerstone of TB treatment. Other studies have shown the interaction between RIF and INH against *M. tuberculosis* to have no interaction or to be mildly antagonistic (8, 65). The inclusion of SPT in this drug regimen was underpinned by reports that 24 out of 70 random combinations tested were synergistically active in *M. smegmatis* (10). This suggests a large unexplored pool of synergistic combinations. Notably, SPT exhibited synergy with several compounds both *in vitro* and *ex vivo* (10), even though the compound has high MIC_90_ against *M. tuberculosis* when administered on its own (17).

In the three-drug combination assay, synergy resulted when sub-inhibitory concentrations (1/2× and 1/4× MICs) of SPT were titrated into media containing RIF and INH. This finding correlated well with the results of time-kill kinetics. However, the time-kill assay suggested that the inhibitory effect of the three-drug interaction was bacteriostatic (0≥ log_10_ CFU <3 reduction) and not bactericidal. This observation reveals that the combination of RIF and INH — two bactericidal drugs that are most potent against actively dividing cells — shows bacteriostatic effects. Furthermore, the inhibition of growth induced by a bacteriostatic drug, SPT, results in an overall static effect when the drug is used in combination with a bactericidal drug. Other studies have shown that, in similar interactions, the resulting effect achieves a more efficient clearance at lower concentrations (40).

In attempting to exploit synergy for potential optimal treatment outcomes, an investigation of the RIF-INH plus SPT interaction was performed in *rpoB* and *inhA* mutants. The RIF^R^ *rpoB* mutant had an MIC value >2000 times the MIC_90_ for drug-susceptible *M. tuberculosis*. Notably, addition of 1/2× MIC of SPT restored partial drug efficacy against this resistant mutant. As with the drug susceptible H37Rv strain, a mechanistic explanation for the synergy observed using the RIF-INH plus SPT combination against the *rpoB* mutant is presently lacking. INH targets mycobacterial cell envelope biosynthesis, possibly enhancing permeation of SPT into the bacilli. However, access alone may not necessarily contribute to the synergistic interaction. Chen *et al*. reported synergy between SQ109, a presumed cell envelope inhibitor, and RIF (34). On the other hand EMB, which also affects mycobacterial cell envelope synthesis, did not exhibit synergy with RIF (66).

Prior reports have shown RIF to be an efficient inducer of cytochrome P450 (CYP 450), a superfamily of haem-containing enzymes involved in the biosynthesis of compounds such as sterols, steroids, and fatty acids as well as detoxification of xenobiotics and chemicals (67). RIF has been linked with the induction of CYP both in humans and in *M. tuberculosis* (68). The elevated levels of CYP have been associated with drug resistance due to the enhanced rate of elimination of the drugs by metabolism and detoxification pathways. INH, on the other hand, inhibits CYP in *M. tuberculosis* (68). This ability of INH to inhibit CYP may contribute to synergy in the RIF-INH plus SPT combination when the active form of INH is not rapidly eliminated inside *M. tuberculosis* and when SPT acts by further reducing the activiy of CYPs.

There are prospects to combine SPT with RIF-INH in treatment regimens. SPT is given by intramuscular injection to achieve therapeutic concentrations in serum of about 100 mg/L one hour after a single 2 g dose. An over 4-fold increase in its effectiveness within a triple SPT-RIF-INH combination, as indicated by these data, would potentially allow for oral formulation, a critical delivery format when administering treatment to TB out-patients. In summary, these *in vitro* and *ex vivo* results suggests that the RIF-INH plus SPT triple-combination may be an effective therapeutic option for the treatment of both the drug susceptible and resistant *M. tuberculosis* infections. They also reinforce a growing body of evidence supporting the utility of drug potentiation strategies in improving treatment outcomes.

## ACKNOWLEDGEMENTS

Research reported in this publication was supported by the Strategic Health Innovation Partnerships (SHIP) Unit of the South African Medical Research Council (SAMRC) with funds received from the South African Department of Science Innovation (to KC and DFW). We acknowledge the support of the University of Cape Town, the SAMRC (to KC and VM), the South African Research Chairs Initiative (to KC) and Department of Science Innovation administered through the National Research Foundation, the Schlumberger Foundation Faculty for the Future scheme (to EK and AW), and the Research Council of Norway (to DFW). We thank Prof. J. A. Aínsa for providing the *M. tuberculosis* Δ*Rv1258c* (“tap”) knockout mutant and its complemented derivative, Prof. R. E. Lee for providing stocks of the spectinamide, 1599, Dr. Krupa Naran for advice with checkerboard assays and Dr. Mandy Mason for critical review of the manuscript.

## TRANSPARENCY DECLARATIONS

None to declare.

