## Supplemental Info for "Developing synergistic drug combinations to restore antibiotic sensitivity in drug-resistant *Mycobacterium tuberculosis*"

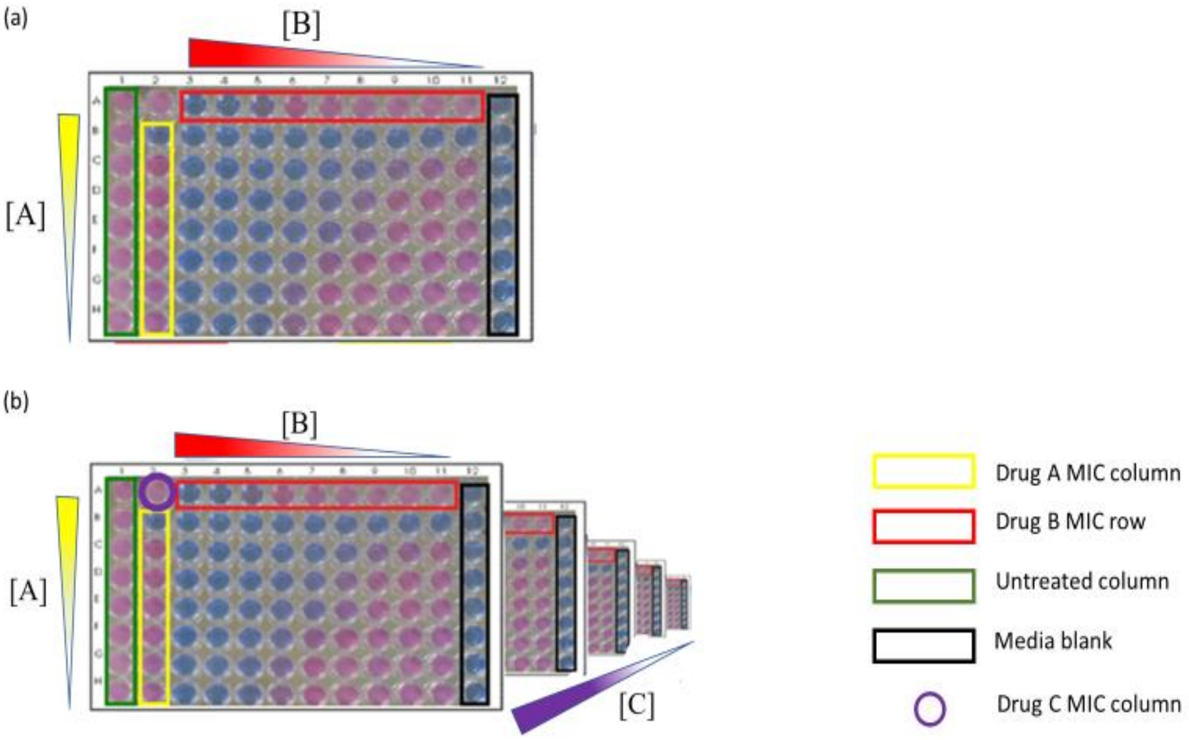


**Figure S1.** Schematic representation of (a) 2D and (b) 3D checkerboard 96-well plate assays. In each assay, the MICs of individual drugs are determined in the "Drug MIC" row/column. The lowest sum FIC (ΣFIC) is calculated from wells containing the combined drugs at different concentrations. For the 3D assay, a third drug [C], is added as an overlay at sub-MIC (five concentrations from 1/2× to 1/32×) on each of the five 96-well plates in the stack.

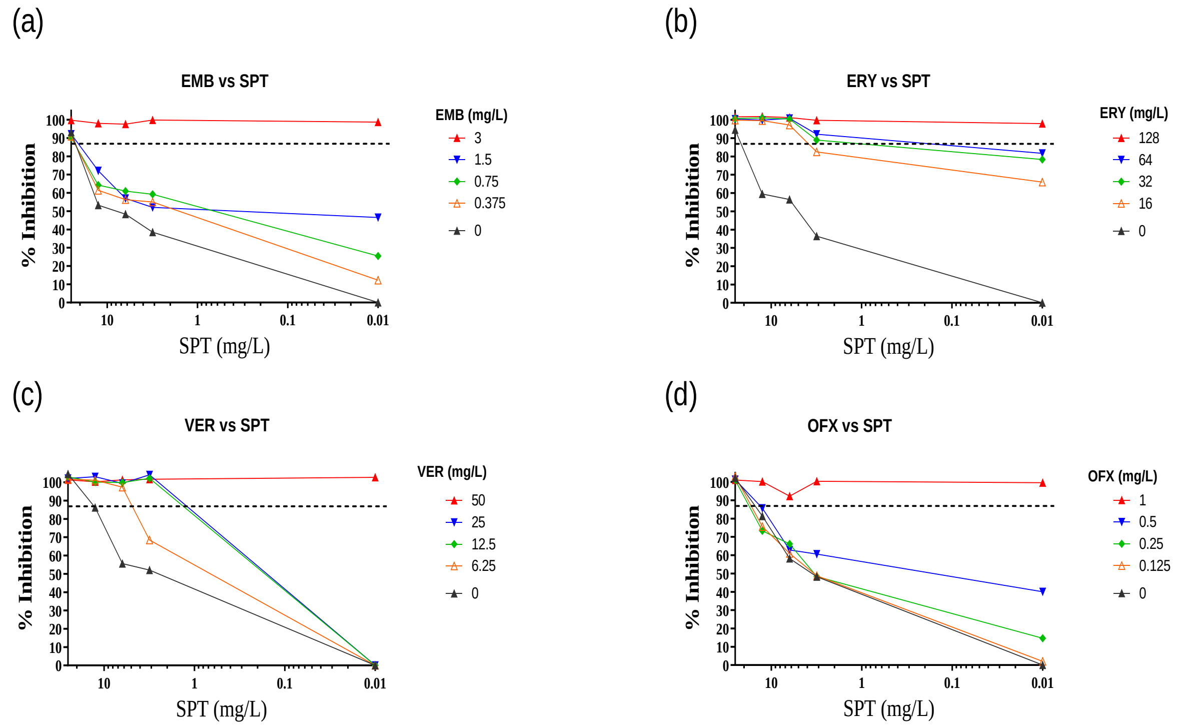

(a)
EMB vs SPT
% Inhibition
SPT (mg/L)
EMB (mg/L)
3
1.5
0.75
0.375
0
(b)
ERY vs SPT
% Inhibition
SPT (mg/L)
ERY (mg/L)
128
64
32
16
0
(c)
VER vs SPT
% Inhibition
SPT (mg/L)
VER (mg/L)
50
25
12.5
6.25
0
(d)
OFX vs SPT
% Inhibition
SPT (mg/L)
OFX (mg/L)
1
0.5
0.25
0.125
0


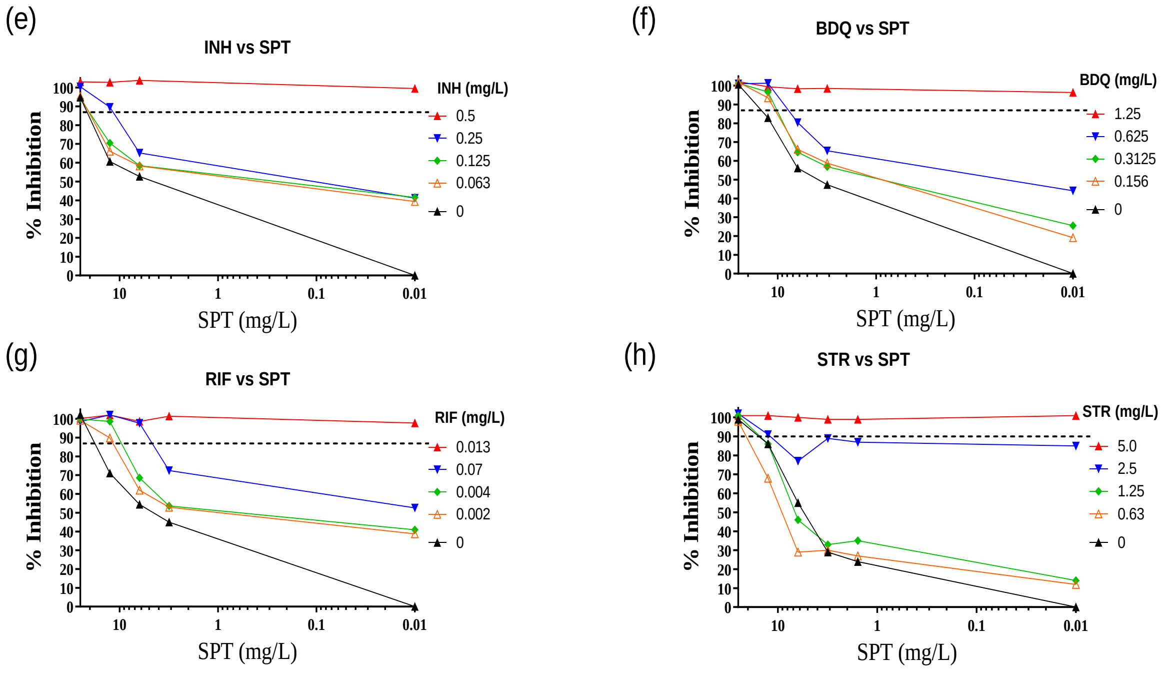

(e)
INH vs SPT
% Inhibition
SPT (mg/L)
INH (mg/L)
0.5
0.25
0.125
0.063
0
(f)
BDQ vs SPT
% Inhibition
SPT (mg/L)
BDQ (mg/L)
1.25
0.625
0.3125
0.156
0
(g)
RIF vs SPT
% Inhibition
SPT (mg/L)
RIF (mg/L)
0.013
0.07
0.004
0.002
0
(h)
STR vs SPT
% Inhibition
SPT (mg/L)
STR (mg/L)
5.0
2.5
1.25
0.63
0

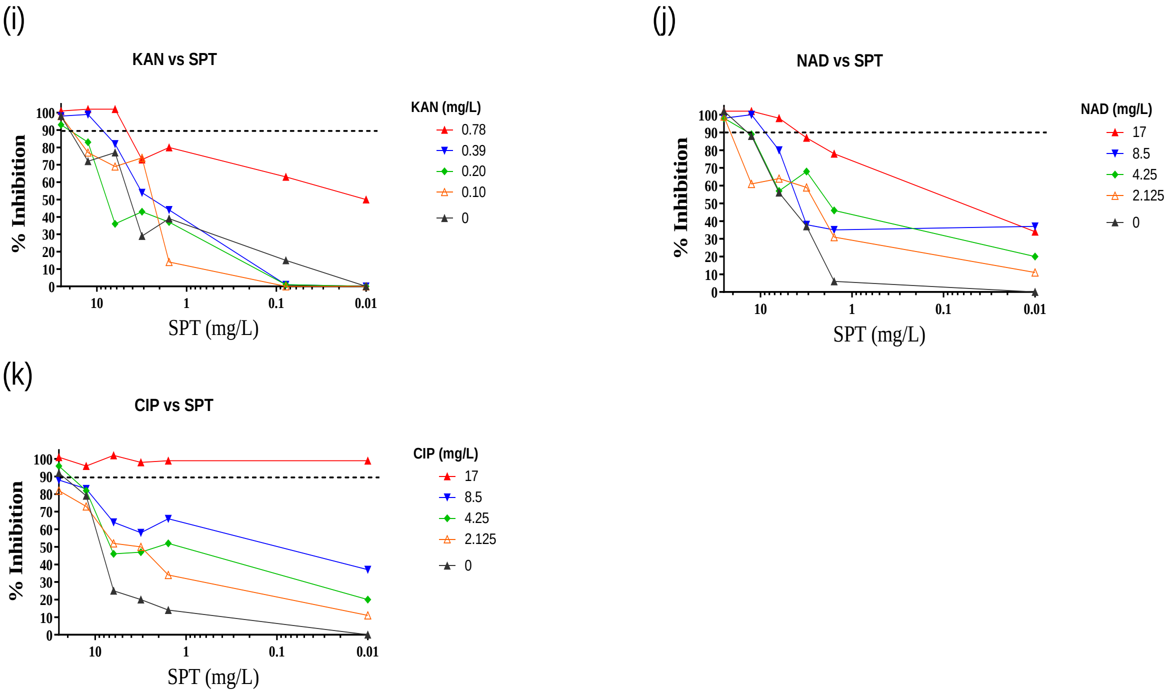


**Figure S2.** *In vitro* interactions between SPT and selected compounds against wild-type *M. tuberculosis* H37Rv. Bacterial viability was assessed in two independent experiments by fluorescence-based resazurin assay. Dashed horizontal lines indicate 90% inhibition. BDQ, bedaquiline; CIP, ciprofloxacin; EMB, ethambutol; ERY, erythromycin; INH, isoniazid; KAN, kanamycin; NAD, nalidixic acid; OFX, ofloxacin; RIF, rifampicin; SPT, spectinomycin; STR, streptomycin; VER, verapamil.

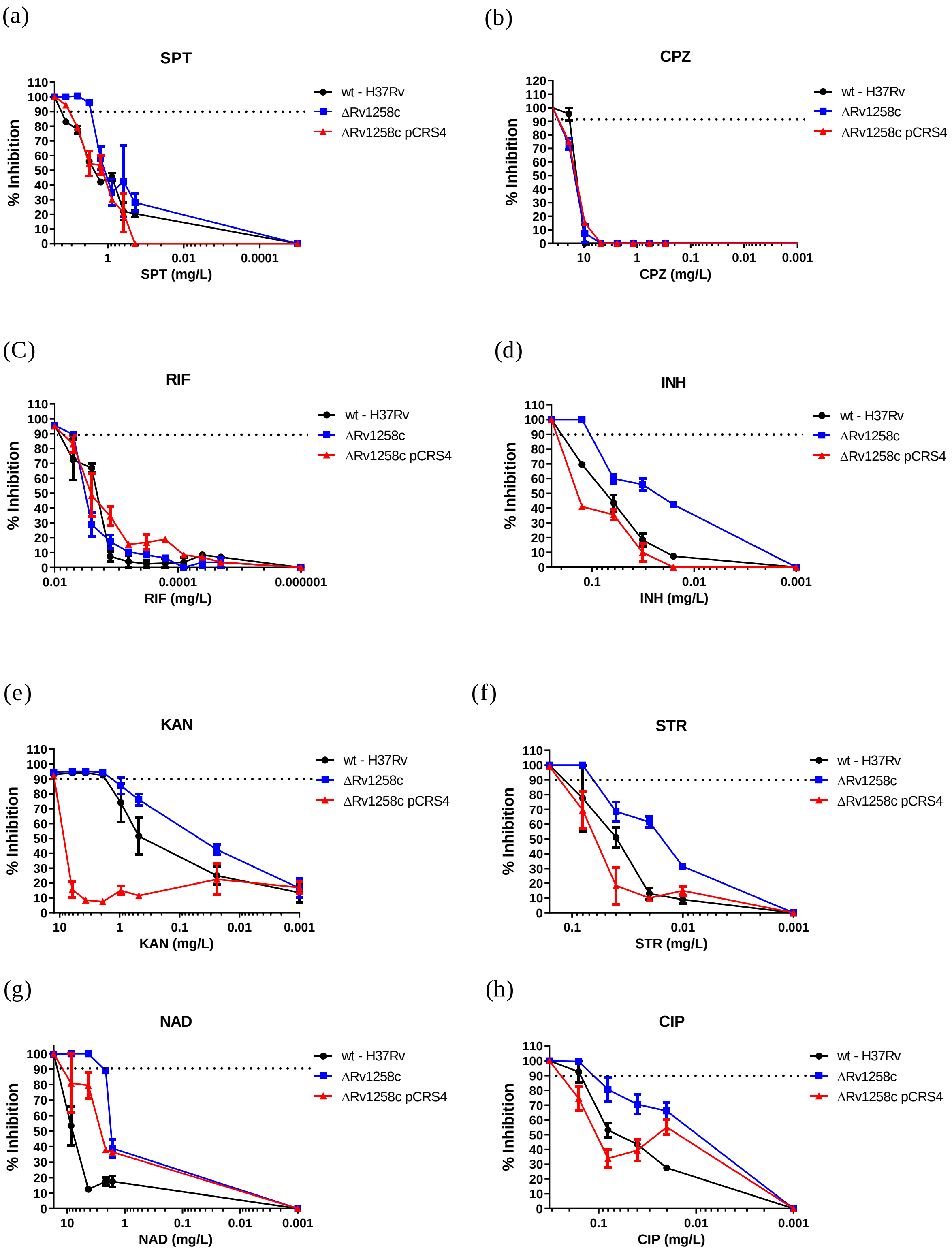
(a)
SPT
% Inhibition
SPT (mg/L)
(b)
CPZ
% Inhibition
CPZ (mg/L)
(C)
RIF
% Inhibition
RIF (mg/L)
(d)
INH
% Inhibition
INH (mg/L)
(e)
KAN
% Inhibition
KAN (mg/L)
(f)
STR
% Inhibition
STR (mg/L)
(g)
NAD
% Inhibition
NAD (mg/L)
(h)
CIP
% Inhibition
CIP (mg/L)
wt - H37Rv
ΔRv1258c
ΔRv1258c pCRS4

(h)

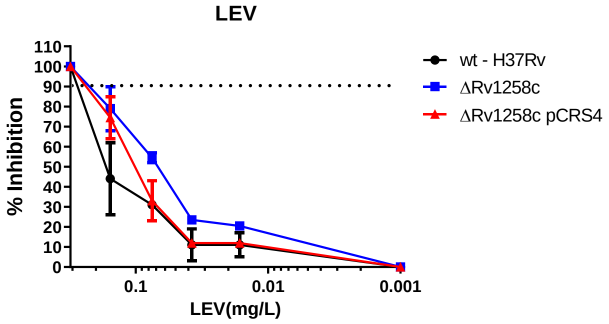


**Figure S3.** Investigating potential hypersensitivity of the *M. tuberculosis* Δ*Rv1258c* mutant to different anti-TB compounds. All compounds were tested against wild-type (H37Rv), the "tap" knock-out (Δ*Rv1258c*), and its complemented derivative (Δ*Rv1258c* pCRS4) on 96-well plates. Bacterial viability was assessed in two independent experiments by fluorescence-based resazurin assay. Dashed horizontal lines indicate 90% inhibition. CIP, ciprofloxacin; INH, isoniazid; KAN, kanamycin; NAD, nalidixic acid; OFX, ofloxacin; RIF, rifampicin; SPT, spectinomycin; STR, streptomycin.

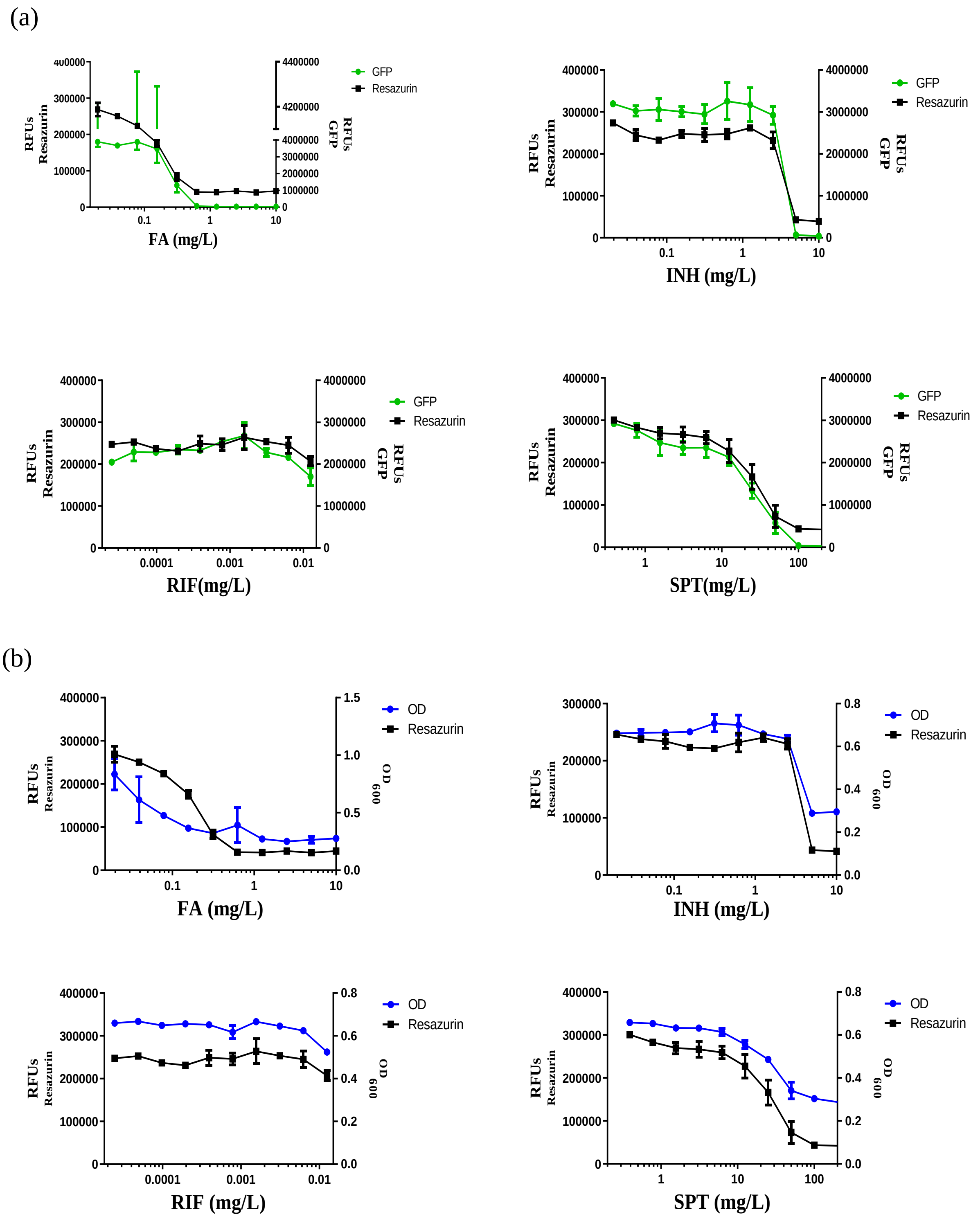


**Figure S4.** Relative fluorescence units (RFU) of the Resazurin assay or GFP assay and absorbance ($OD_{600}$) in calculating MIC values against wild-type *M. tuberculosis* H37Rv. (a) Resazurin *versus* GFP fluorescence intensities; (b) resazurin fluorescence intensities *versus* $OD_{600nm}$ readings.

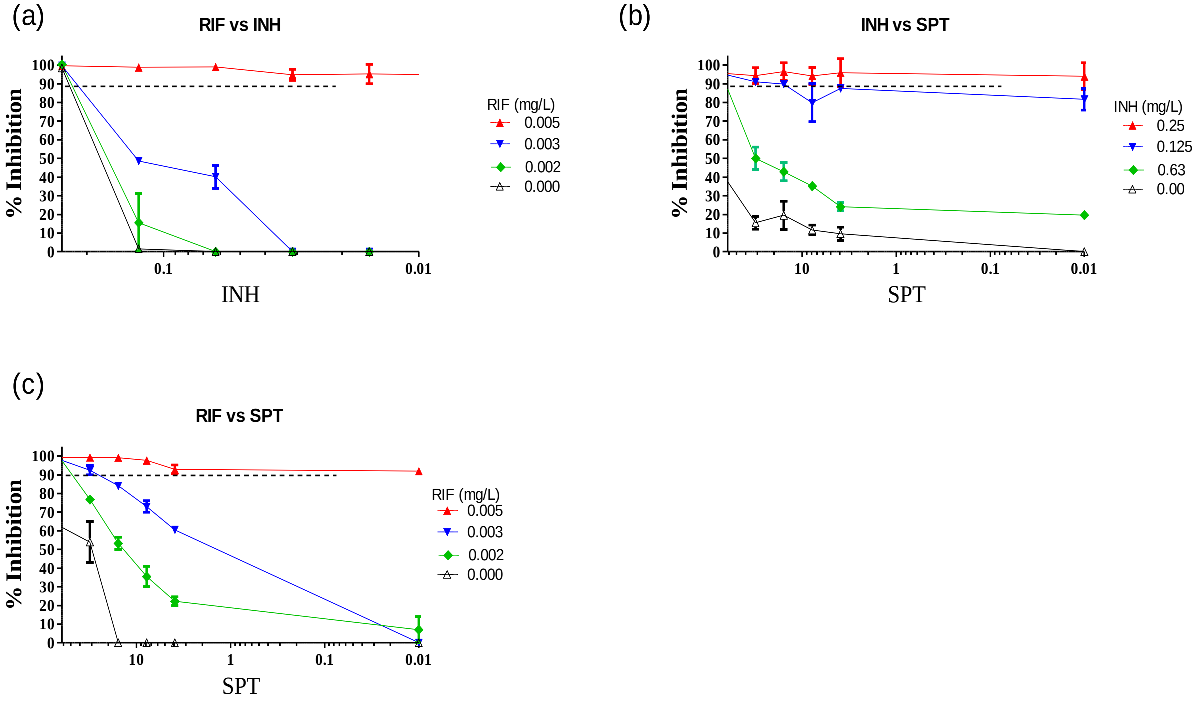


**Figure S5.** *In vitro* interactions between (a) RIF and INH, (b) INH and SPT, and (c) RIF and SPT in 2D checkerboard assays against wild-type *M. tuberculosis* H37Rv. Bacterial viability was assessed in two independent experiments by fluorescence-based resazurin assay. Dashed horizontal lines indicate 90% inhibition.

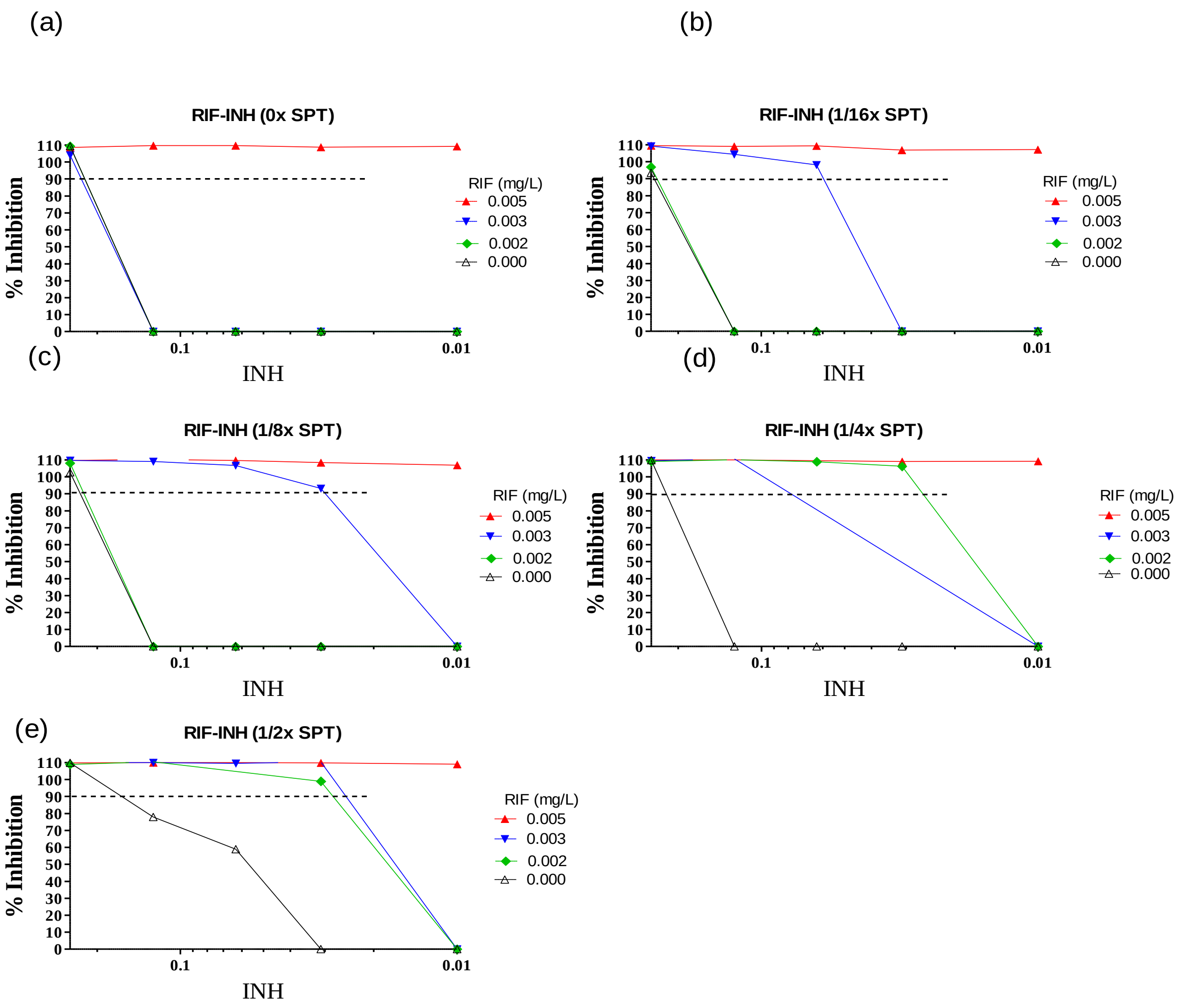


**Figure S6.** 3D checkerboard assay of RIF and INH against wild-type *M. tuberculosis* H37Rv in the presence of sub-$MIC_{90}$ concentrations of SPT: (a) 0x SPT, (b) 1/16× SPT, (c) 1/8× SPT, (d) 1/4× SPT, and (e) 1/2× SPT. Bacterial viability was assessed in two independent experiments by fluorescence-based resazurin assay. Dashed horizontal lines indicate 90% inhibition.

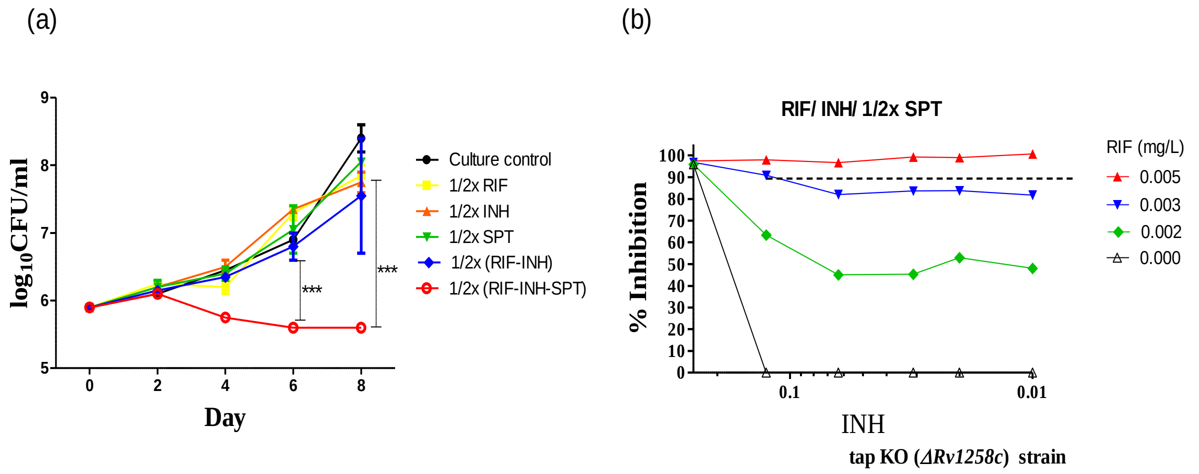


**Figure S7.** *In vitro* interactions between RIF and INH in the presence of a sub-inhibitory concentration of SPT (1/2× MIC SPT). (*a*) Kill kinetics determined following exposure of wild-type *M. tuberculosis* H37Rv to sub-inhibitory concentrations of RIF, INH and SPT individually and in combination. Viable cell counts were determined at the indicated timepoints by plating and enumerating colony forming units (CFU) on antibiotic-free Middlebrook 7H10 agar ($p < 0.001$). (*b*) Checkerboard assays were performed using RIF, INH and 1/2×MIC SPT against the Δ*Rv1258c* mutant ("tap KO strain") in a 96-well microtiter plate. The fluorescence reading was obtained on Day 8 following the addition of resazurin. Data are representative of two independent biological replicates.

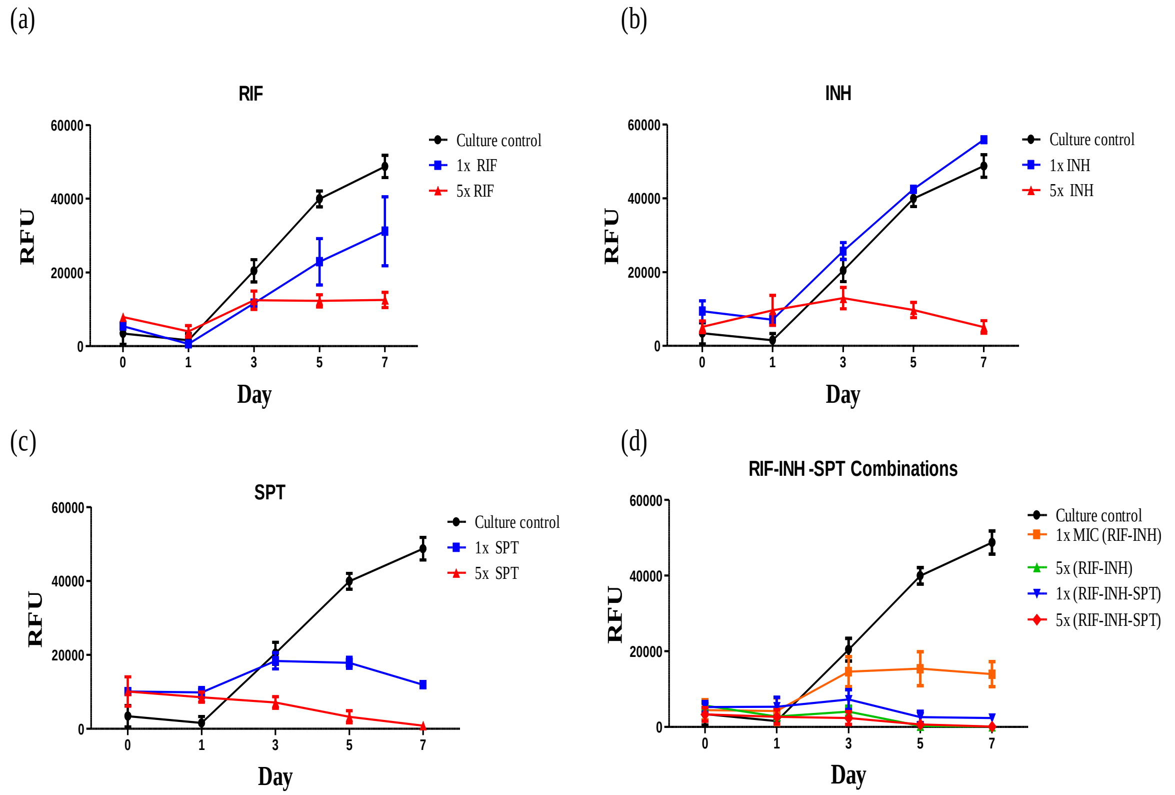


**Figure S8.** Activities of RIF, INH and SPT alone and in combination in a model of *M. tuberculosis* intracellular infection. THP-1 cells were infected with exponentially growing *M. tuberculosis* H37Rv pSMYC::mCherry) at a multiplicity of infection (MOI) of 1:5 cells:bacilli. Following infection, cells were exposed to different concentrations of each of the partner drugs alone or in combination. The plots detail the inhibitory activities of the drugs as determined by mCherry fluorescence intensity. Data are from independent experiments performed in triplicate.

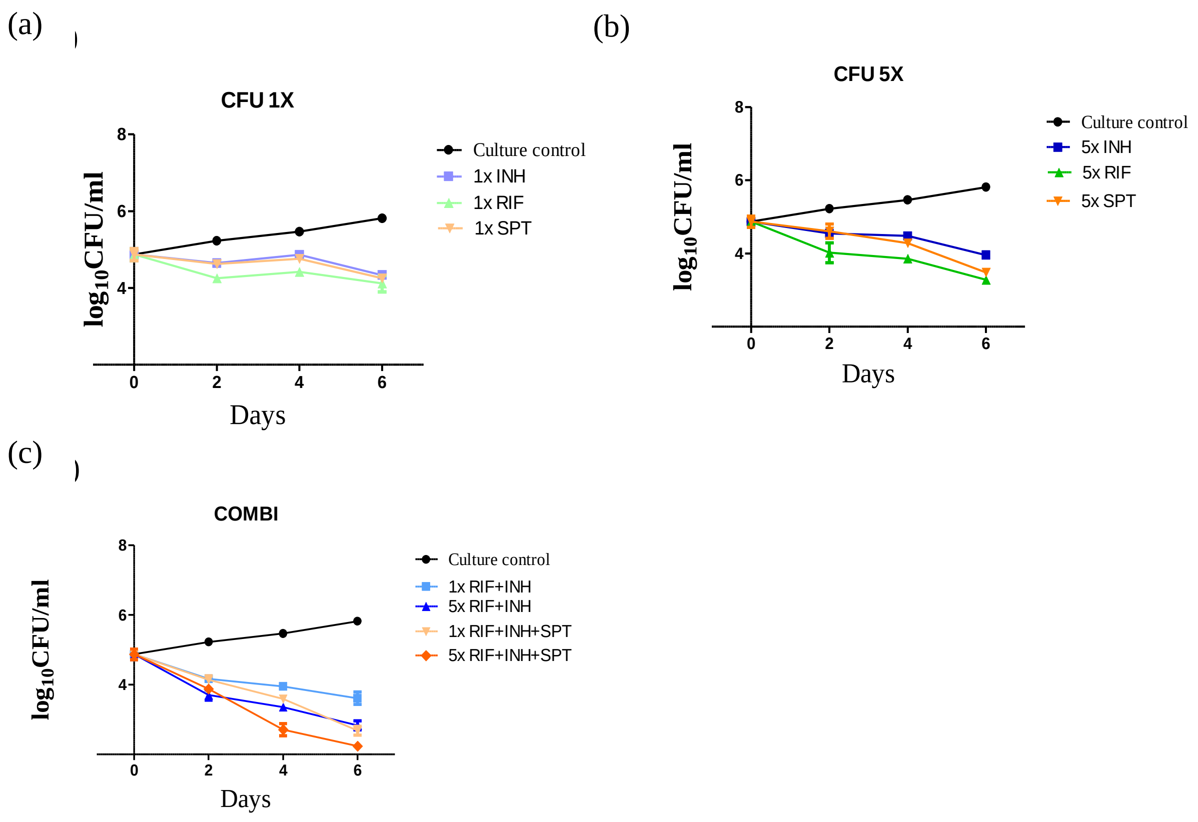


**Figure S9.** Intracellular activities of RIF, INH and SPT alone and in combination. THP-1 cells were infected with *M. tuberculosis* H37Rv at a multiplicity of infection (MOI) of 1:5 cells:bacilli. Following infection, cells were exposed to different concentrations of each of the partner drugs alone (“CFU 1X”, “CFU 5X”) or in combination (“COMBI”). The plots detail the inhibitory activities of the drugs as determined by CFU enumeration following plating on antibiotic-free Middlebrook 7H10 solid medium. Data are from independent experiments performed in triplicate.

**Table S1** *In vitro* activities of selected compounds against wild-type *M. tuberculosis* H37Rv

| | | | | $MIC_{90}$ (mg/L) | | | |
|---|---|---|---|---|---|---|---|
| **Compound** | **H37Rv** | **FA$^R$** | **SPT$^R$** | **Δ*Rv1258c*†** | **Δ*Rv1258c* pCRS4** | ***inhA*** | ***rpoB*** |
| FA | 0.63 | >248 | 0.63 | 0.32 | 0.63 | | |
| SPT | 62-125 | 125 | >8000 | **3.9** | 31 | | 62 |
| ‘1599 | 3.12 | | | 3.12-1.56 | 6.25 | | |
| CPZ | 22.2 | 22.2 | 22.2 | 22.2 | 22.2 | | |
| RIF | 0.01 | | 0.01 | 0.005 | 0.01 | 0.01 | 20 |
| INH | 0.25 | | 0.125 | 0.125 | 0.25 | >4.0 | 0.06 |
| KAN | 1.56 | | | 1.56 | 12.5 | | |
| LEV | 0.31 | | | 0.16-0.31 | 0.31 | | |
| NAD | 17 | | | 8.5-17 | 17-34 | | |
| CIP | 0.16 | | | 0.16 | 0.32 | | |
| STR | 0.08 | | | 0.08 | 0.08-0.16 | | |
| BDQ | 1.2 | | | 0.6 | 1.2 | | |

†$MIC_{90}$ values in bold type indicate hypersensitivity of the Δ*Rv1258c* mutant to the specific compound(s)

**Table S2:** $MIC_{90}$ and ΣFIC values for selected antimycobacterial compounds used against wild-type *M. tuberculosis* H37Rv alone and in combination with CPZ

| **Compound** | **$MIC_{90}$ (mg/L), alone** | **$MIC_{90}$ (mg/L), in combination** | **FIC** | **ΣFIC†** |
|---|---|---|---|---|
| RIF | 0.0032 | 0.00037 | 0.116 | **0.37** |
| CPZ | 22 | 5.50 | 0.250 | |
| INH | 0.38 | 0.094 | 0.247 | **0.50** |
| CPZ | 22 | 5.50 | 0.250 | |
| KAN | 7.16 | 3.57 | 0.499 | 1.00 |
| CPZ | 22 | 11 | 0.500 | |
| STR | 1.15 | 0.14 | 0.122 | 0.62 |
| CPZ | 22 | 11 | 0.500 | |
| BDQ | 3.00 | 0.0059 | 0.002 | **0.25** |
| CPZ | 11 | 2.25 | 0.250 | |
| NAD | 14 | 1.8 | 0.125 | **0.25** |
| CPZ | 22 | 2.75 | 0.125 | |
| CIP | 2.50 | 1.27 | 0.508 | 0.76 |
| CPZ | 22 | 5.50 | 0.250 | |
| LEV | 2.30 | 2.30 | 1.00 | 2.00 |
| CPZ | 22 | 22 | 1.00 | |

†Bold text indicates synergistic combinations (ΣFIC≤0.5)

**Table S3:** $MIC_{90}$ and ΣFIC values for selected antimycobacterial compounds used against wild-type *M. tuberculosis* H37Rv alone and in combination with SPT

| Drugs/Compound | $MIC_{90}$ (mg/L), singly | $MIC_{90}$ (mg/L) in combination | FIC | ΣFIC† |
|---|---|---|---|---|
| RIF | 0.013 | 0.002 | 0.015 | 0.52 |
| SPT | 25 | 12.5 | 0.500 | |
| INH | 0.5 | 0.25 | 0.500 | 1.00 |
| SPT | 25 | 12.5 | 0.500 | |
| BDQ | 1.25 | 0.156 | 0.125 | 0.63 |
| SPT | 25 | 12.5 | 0.500 | |
| OFX | 1.0 | 0.5 | 0.500 | 1.50 |
| SPT | 25 | 25 | 1.000 | |
| EMB | 3.00 | 1.5 | 0.500 | 1.50 |
| SPT | 25 | 25 | 1.000 | |
| ERY | 128 | 32 | 0.125 | **0.38** |
| SPT | 25 | 6.25 | 0.250 | |
| VER | 50 | 6.25 | 0.125 | **0.25** |
| SPT | 25 | 6.25 | 0.125 | |
| KAN | 1.56 | 1.56 | 1.000 | 1.25 |
| SPT | 25 | 6.125 | 0.250 | |
| STR | 0.08 | 0.16 | 2.000 | 2.50 |
| SPT | 25 | 12.5 | 0.500 | |
| CIP | 0.16 | 0.32 | 2.000 | 2.25 |
| SPT | 25 | 6.25 | 0.250 | |
| NAD | 17 | 34 | 2.000 | 2.13 |
| SPT | 25 | 6.25 | 0.125 | |

†Bold text indicates synergistic combinations (ΣFIC≤0.5)
